# A Plausible Accelerating Function of Intermediate States in Cancer Metastasis

**DOI:** 10.1101/828343

**Authors:** Hanah Goetz, Juan R. Melendez-Alvarez, Luonan Chen, Xiao-Jun Tian

**Affiliations:** School of Biological and Health Systems Engineering, Arizona State University; Shanghai Institutes for Biological Sciences, Chinese Academy of Sciences; Center for Excellence in Animal Evolution and Genetics, Chinese Academy of Sciences

**Author notes:** These authors contributed equally to this work (Hanah Goetz, Juan R. Melendez-Alvarez).

**Keywords:** Cancer cell plasticity, partial EMT, microstate, macrostate, intermediate state, Markov process, stabilized state

## Abstract

Epithelial-to-mesenchymal transition (EMT) is a fundamental cellular process and plays an essential role in development, tissue regeneration, and cancer metastasis. Interestingly, EMT is not a binary process but instead proceeds with multiple partial intermediate states. However, the functions of these intermediate states are not fully understood. Here, we focus on a general question about how the number of partial EMT states affects cell transformation. First, by fitting a hidden Markov model of EMT with experimental data, we propose a statistical mechanism for EMT in which many unobservable microstates may exist within one of the observable macrostates. Furthermore, we find that increasing the number of intermediate states can accelerate the EMT process and that adding parallel paths or transition layers accelerates the process even further. Last, a stabilized intermediate state traps cells in one partial EMT state. This work advances our understanding of the dynamics and functions of EMT plasticity during cancer metastasis.

## Introduction

Epithelial-to-mesenchymal transition (EMT) is a fundamental cellular process in which polarized epithelial cells lose various cell-cell junctions and adhesion and gain migratory and invasive properties to become mesenchymal cells [1, 2]. EMT is very important in embryonic development, tumorigenesis, metastasis, tumour stemness, and therapy resistance [3, 4]. Remarkably, EMT is not a binary process but proceeds with multiple partial intermediate states, collectively known as partial or hybrid EMT states [3, 5–11]. The partial EMT state retains some characteristics of epithelium but also shows features of mesenchymal cells [12–14]. One partial EMT state was predicted through mathematical modeling of the EMT core regulatory network and was verified with quantitative experiments by our previous works [5, 6]. Thereafter, many different partial EMT states were proposed [8, 9, 15–17]. More and more experimental data shows a different number of partial EMT states in various cancer cell lines [18–23]. Recently, several partial EMT phenotypes were found during cancer metastasis *in vivo* in a skin cancer mouse model [24, 25] and prostate cancer [26]. While many partial EMT states have been found, their functions are still not fully understood during cancer metastasis [4, 27–2 9].

Currently, the function of partial EMT states has being studied in the context of coupling with other cellular processes. For example, acquisition of stem-like properties dictates its coupling with cancer stemness [11, 30–34], circulating tumor cells (CTCs) [35, 36], and drug resistance [37]. Thus, the partial EMT cells hold the highest metastatic potential. As well, partial EMT, instead of full EMT, is found to be critical for renal fibrosis [38–40]. There are many potential couplings of partial EMT and other biological processes, such as cell cycle [40], renal fibrosis [41] and metabolisms [42]. The cross-talk among the regulators of EMT and other cellular processes guides the coupling mechanism and full functional characteristics of partial EMT states.

While it is important to investigate specific functions of each partial EMT state in the context of additional cellular processes, an interesting question about the general function is whether the number of partial EMT states affect the transition itself. Given that different cell lines may have different partial EMT states, epithelial cells in different systems may also undergo different steps to become mesenchymal cells. Here, we use a general hidden Markov model to describe cell fate transitions during the EMT process. Our analysis makes several non-intuitive predictions. First, several microstates exist within one macrostate, which makes EMT a non-Markov process. Second, increasing the number of intermediate states can accelerate the EMT process. Third, transition in parallel and layer modes can further accelerate the process. Lastly, the existence of a stabilized intermediate state traps cells within their current phenotype. We conclude with discussions on how the dynamics and functions of EMT plasticity are controlled by the number of states during cancer metastasis.

## Results

### EMT is a Non-Markov Process

Three steady states were found in TGF-*β* induced EMT in the MCF10A cell line with quantitative measurements of E-cadherin and Vimentin [6], consistent with theoretical prediction [5]. That is, EMT progresses through three functional cell states in this cell line: from the initial epithelial state to the partial EMT state, where cells lose some of the cell-cell adhesion, and then to the full mesenchymal state (Fig. 1A). As shown in Fig. 1B (left), this system can be represented with a metaphorical three-well landscape in one dimension along the mesenchymal marker (M-Marker) axis. Each well represents a stable cell phenotype and the transitions among them are indicated with arrows. With high concentration of TGF-*β*, the cell will be driven from the epithelial state (the first well) to the mesenchymal state (last well) through the partial EMT state (middle well), indicated by solid arrows. The dynamics of the M-Marker (measured by the Vimentin protein) at the single cell level shows two steps (Fig. 1B, right) [5].

**Fig 1.**
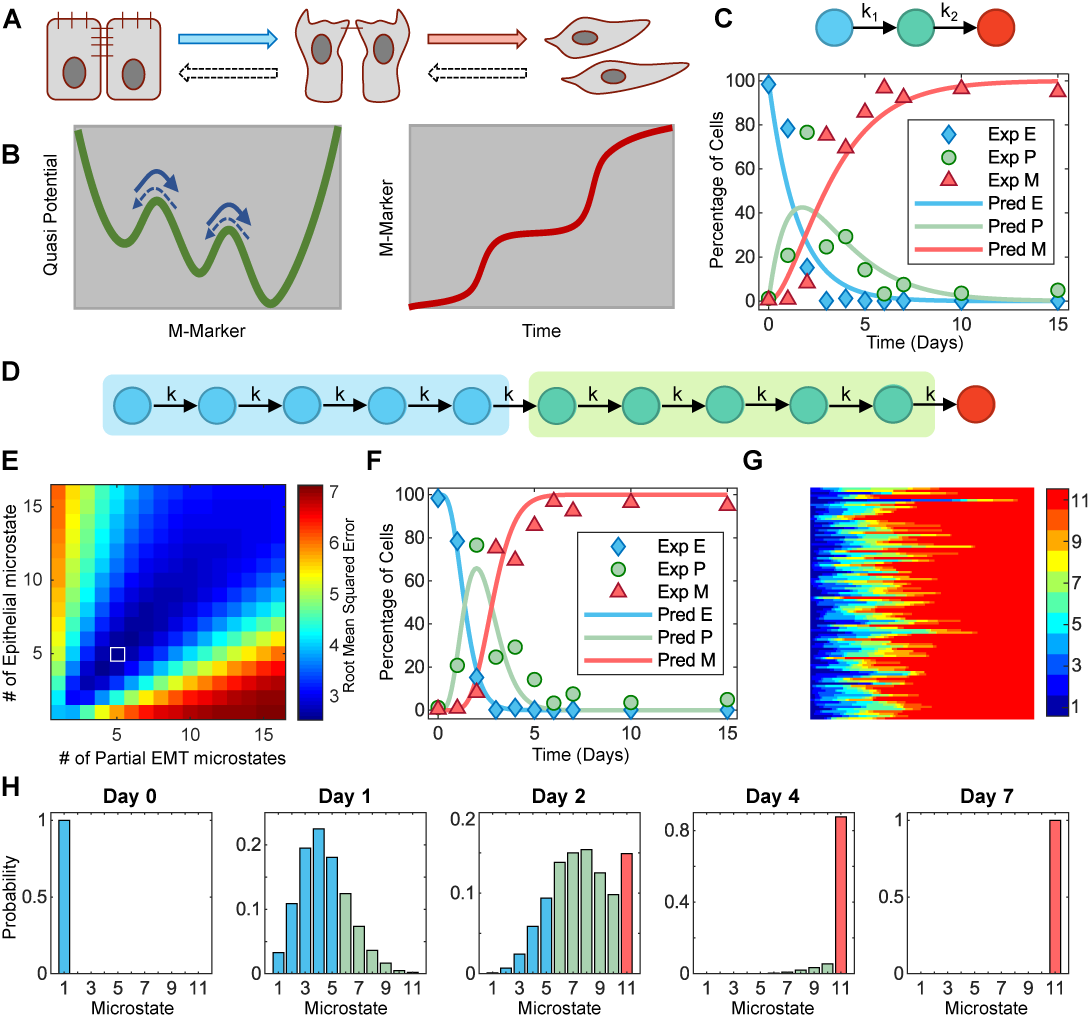
EMT is a non-Markov Process. (A) Cell phenotype transitions among epithelial, partial EMT, and mesenchymal states during EMT. (B) Metaphorical three-well landscape in one dimension along the mesenchymal marker (M-Marker) axis shows the cell phenotype transition during EMT. Full arrows show the order of EMT while dashed arrows show reverse process, Mesenchymal-to-Epithelial transition (MET). (C) Three-state Markov model for EMT. Best fit of the model to previous experimental data on temporal changes of the percentage of cells in three states during EMT [6] with *k*_1_ = 0.6657 and *k*_2_ = 0.4908 (bottom). (D) An extended model with *N*_*E*_ epithelial microstates (blue circles) and *N*_*P*_ partial EMT microstates (green circles). (E) The fitting score, root mean squared error (RMSE), with the extended model in the space of *N*_*E*_ and *N*_*P*_. The best fit is at *N*_*E*_ = 5 and *N*_*P*_ = 5 (white square, RMSE=2.55, k=3.4261). (F) The dynamics of cells in three macrostates during EMT with the best-fitted extended model overlaid with experimental data. (G) Stochastic simulation shows the state transition of 100 simulated cells over time. (H) Probability of cells in each microstate at several time points. 10,000 simulated cells with a stochastic model (see the methods for details) are sampled to calculate the probability.

Due to the cell-cell variability, the cells do not synchronize the transition through these states but make the transitions in a stochastic manner. However, the fraction of cells in each state show deterministic dynamics, as shown by the quantified experimental data (Fig. 1C) from Ref. [6]. To better understand EMT at the population level, we first build a simple model by assuming that cells are initially held in the epithelial state and will be driven by a high dose of TGF-*β* as used in the experiments to the partial EMT state before ending in the mesenchymal state, all with constant average rates (*k*_1_ and *k*_2_) (Fig. 1C). We fit this simple model with the experimental data and find that the model does not describe the data with strong accuracy (Fig. 1C). With this model, the transition from the partial EMT state to the mesenchymal state shows a quicker dynamic than the observed data. Thus, this model is not a good representation of the EMT process at the population level.

To find out the underlying mechanism for the observed dynamics, we extend the simple model by adding several microstates for each cell macrostate (Fig. 1D) by assuming that some unmeasured variables can further distinguish the epithelial state and partial EMT state into several microstates, with the epithelial and partial EMT states being the macrostates used in describing the overall EMT process. While we do not know the actual number of the epithelial and partial EMT microstates (*N*_*E*_ and *N*_*P*_, respectively), a fitting of the extended model is done with all the combinations of *N*_*E*_, *N*_*P*_, and *k* (Fig. 1E) by assuming an equal transition rate from one state to the following state (*k*). According the fitting score (root mean squared error), the best fit is located when *N*_*E*_ = 5 and *N*_*P*_ = 5 (square in Fig. 1E). The best fitting shown in Fig. 1F shows the optimal conditions in which our model reproduces the EMT dynamics at the population level with better accuracy than that of the assumption of only one microstate in both the epithelial and partial EMT states. These results suggest that EMT is a non-Markov process with many hidden epithelial microstates and partial EMT microstates, which could further be distinguished by measuring other variables in the system.

This conclusion is further confirmed with a stochastic model to demonstrate the evolution of cell transitions in these microstates (see the method for details). Fig. 1G shows 100 typical simulated stochastic trajectories of cells to complete the EMT process. With all cells beginning in the epithelial state, prominent cell-to-cell variation is shown in the time it takes a cell to convert to different microstates, indicating unsynchronized transition at the single cell level. At the population level, the statistic results with 10,000 simulated cells shows the dynamics of the distribution of cells in the microstates (Fig. 1H). Few reach the mesenchymal state (red bar) by day two. However, most cells will reach the final EMT state by day four. This is also consistent with the experimental data [6] (Fig. S1A). Furthermore, fitting the experimental data with a model by considering the reversible state transitions suggests more microstates (Fig. S2A). Taken together, our analysis based on fitting a general Markov model with the experimental data suggests that EMT is a non-Markov process, where several microstates exist within the epithelial and partial EMT states.

### Increasing the Number of Intermediate States Can Accelerate EMT

Our analysis with a Markov model for EMT at the population level based on the experimental data of MCF10A cells suggests that multiple microstates exist in each macrostate during the EMT process. However, the number of microstates and macrostates can be cell-type specific. Two questions arise concerning whether and how the number of microstates affect EMT dynamics. For simplification of discussion, we will not distinguish the microstate and macrostates, but instead designate them as intermediate states if they are not either the initial or final state. To compare the systems with different numbers of intermediate states, we made one assumptions that the total energy barrier for the full transition from epithelial (E) to mesenchymal (M) is the same. As shown in Fig. 2A (left), if the energy barrier from E to M is Δ*E* without any intermediate states (blue curve), adding one additional state makes the transition two steps and the barrier is Δ*E/*2 (red curve) for each step. Similarly, the barrier is Δ*E/*3 for each step if two intermediate states are considered (Fig. 2A, right, red curve). This assumption comes from the hypothesized monotonical energy gradient based on epigenetic changes between different EMT states [43] and existence of ‘checkpoints’ in the EMT continuum [44]. This assumption is also consistent with the cascading bistable switches mechanism in which EMT proceeds through step-wise activation of multiple feedback loops [5, 6, 9, 45]. The cells need to change the profiles of gene expression to make the full EMT transition. Under this assumption with one intermediate state, the cells can make changes on the expression of some genes as the first step for a partial transition, and then make changes on the rest of the genes for the second step to complete the transition. The profiles of the genes that control the eukaryotic cell phenotypes are usually sophisticated by positive feedback loops to avoid undesired random phenotype switching from noises or short pulse stimulus. These feedback loops can be mutual inhibitions between two groups, one of them controls cell phenotype one while the other group controls cell phenotype two. Thus, the cell fate transitions through EMT intermediate states are analogous to crossing a series of energy barrier which are controlled by these positive feedback loops.

**Fig 2.**
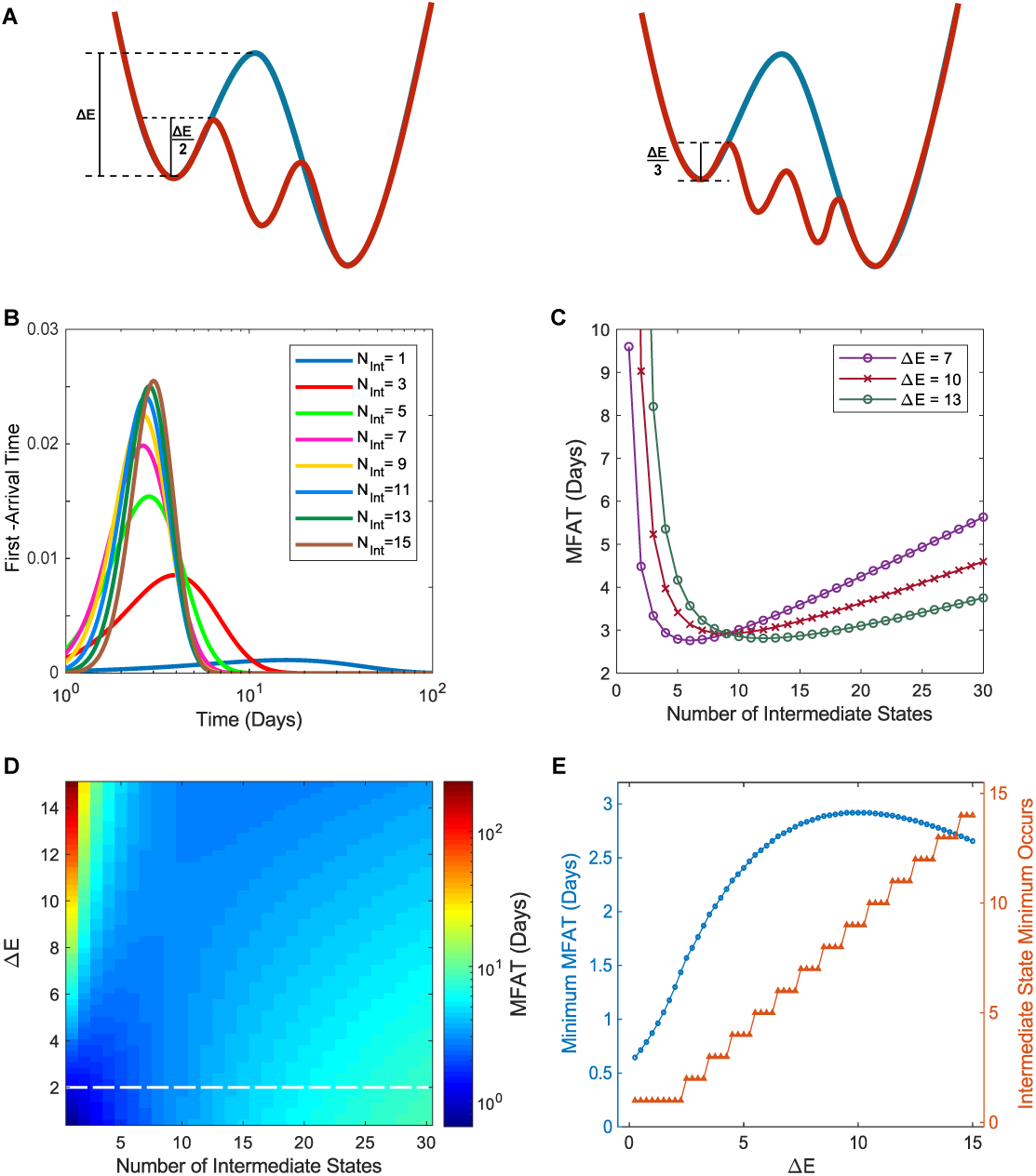
Increasing the number of intermediate states could accelerate the EMT Process. (A) The assumptions that the total energy barrier (Δ*E*) for the full transition from epithelial to mesenchymal is fixed to compare the effects of the number of intermediate states (*N*_*int*_) on the dynamics of EMT. The energy barrier is Δ*E/*(*N*_*int*_ + 1) for each step for the system with *N*_*int*_ intermediate states. (B) The distribution of the first arrival time to the mesenchymal state with different *N*_*int*_ and fixed Δ*E* = 10. (C) The dependence of the mean first arrival time (MFAT) on *N*_*int*_ for three different energy barrier values, as indicated. (D) MFAT in the space of Δ*E* and *N*_*int*_. A dashed white line represents the threshold above which MFAT shows a nonmonotonic dependence on *N*_*int*_. (E) The dependence of the minimum MFAT (blue) on Δ*E* alongside the intermediate state that this minimum occurs (orange).

First, we systematically study how the EMT process changes with fixed Δ*E* and the number of intermediate states, *N*_*int*_. The distributions of the first-arrival time (FAT), defined as the time the cells take to arrive to the mesenchymal state, is used to measure how fast the EMT process is. As shown in Fig. 2B, with increase of *N*_*int*_, the FAT distribution first shifts to the left and then to the right, becoming increasingly more narrow. To see this affect more clearly, we further calculate the mean first arrival time (MFAT) and find a nonmonotonic dependence of MFAT on *N*_*int*_ (Fig. 2C). As *N*_*int*_ increases, the MFAT first shows a rapid decrease. However, after a certain *N*_*int*_, the MFAT begins to slowly increase again. These data suggests that the partial intermediated states have a potential function of accelerating the EMT process. Furthermore, we analyze how Δ*E* affects the EMT process. Shown in Fig. 2C, the MFAT displays the same nonmonotonic pattern for three different Δ*E* values, although the position of the minimum varies. At these minimum points, the EMT process is completed the fastest. This interesting phenomena is further confirmed by the analysis of the MFAT in the space of Δ*E* and *N*_*int*_ (Fig. 2D). The same trend of MFAT can be observed for all Δ*E >* 2 (threshold shown by the dash line). This result suggests that EMT process is not continuous process but proceeds with finite number of discrete microstates.

For the Δ*E* lower than the threshold, MFAT constantly increases with *N*_*int*_ (Fig. 2D). The distribution of the first-arrival time for one case with small Δ*E* is shown in Fig. S1B. One can see that the transition time is already very small in this case and that increasing intermediate states barely increases the transition rate for each step, thus causing more time to complete full EMT. This is largely not the case for the EMT process in a real system, as the minimal MFAT is less than one day, but it demonstrates the underlying mechanism of the nonmonotonic dependence of MFAT on *N*_*int*_. That is, increasing *N*_*int*_ after the minimum does not help to increase the transition rate for each step, it just increases the steps. Thus, there is a trade-off to accelerate EMT through increasing the number of intermediate states. The dependence of the minimum MFAT alongside the intermediate state in which it occurs 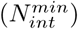 is shown in Fig. 2E. With increase of Δ*E*, both the minimal MFAT and 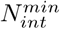 increase, although a small decline of minimal MFAT is observed for large Δ*E*. Taken together, a potential function of multiple partial EMT states is to accelerate EMT, and the number of intermediate states necessary to achieve fastest EMT increases with the total energy barrier.

### Adding Parallel Paths or Transition Layers Further Accelerates the EMT Process

We have discussed the scenario with only one path in which epithelial cells proceed through a step-wise transition of intermediate states to become mesenchymal cells. Here, we consider multiple paths in parallel (Fig. 3A) and study how the number of paths (*N*_*pth*_) affects EMT. For *N*_*int*_ of intermediate states, there are *N*_*int*_ of possible cases (*N*_*pth*_ = 1 ∼*N*_*int*_) based on the one simplification that each path contains close to the same number of intermediate states. For example, if there are nine intermediate states and two paths, four states would belong to one path and five to the other (Fig. 3A).

**Fig 3.**
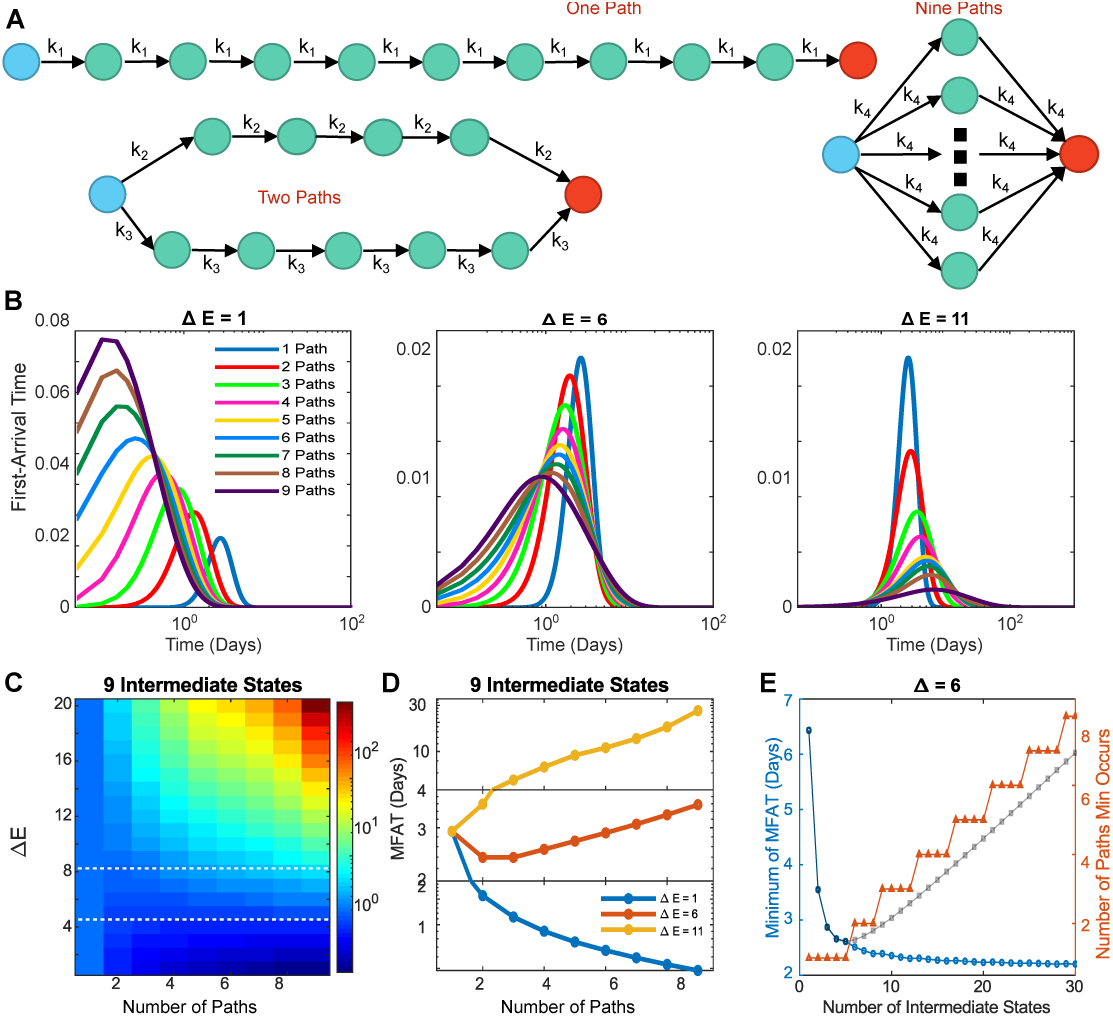
Adding parallel paths can further accelerate the EMT Process. (A) Diagram of cell phenotype transition through nine intermediate states with multiple parallel paths. (B) Dependence of FAT distribution on *N*_*pth*_ with three different Δ*E* values and *N*_*int*_ = 9. (C) Phase diagrams showing MFAT subjected to *N*_*pth*_ and Δ*E* with *N*_*int*_ = 9. Dashed lines represent two thresholds for monotonic and nonmonotonic dependence of MFAT on *N*_*pth*_. (D) Three typical examples show the monotonic and nonmonotonic dependence of MFAT on *N*_*pth*_ according to Δ*E* value. (E) The dependence of minimal MFAT (blue curve) with a comparison to one parallel path (grey dashed line), and the *N*_*pth*_ the minimum occurs on (red) for Δ*E* = 6.

To demonstration the effect of multiple paths, we first study the system with nine intermediate states (*N*_*int*_ = 9). The FAT distribution to the mesenchymal state is shown in Fig. 3B under various *N*_*pth*_ and three levels of energy barrier, Δ*E*. For small Δ*E* (left panel), the FAT distribution shifts to the left as *N*_*pth*_ increases, implying that more paths create a faster transition. However, for large Δ*E* (right panel), the FAT distribution reflects the opposite, implying that more paths create a slower transition. These also suggests a nonmonotonic dependence of FAT on *N*_*pth*_ for moderate Δ*E*. As shown in Fig. 3B (middle panel), the FAT distribution peak shifts to the left but the distribution width becomes larger with increase of *N*_*pth*_. This suggests that the dependence of the FAT on *N*_*pth*_ is monotonic or nonmonotonic according to the energy barrier.

To further confirm this conclusion, we systematically study how the MFAT of the mesenchymal state changes in the space of Δ*E* and *N*_*pth*_ (Fig. 3C). The MFAT dependence on *N*_*pth*_ indeed shows three trends according to the value of Δ*E*. For small (below the bottom white dashed line) or big (above the top white dashed line) values of Δ*E*, the dependence is monotonical, either decreasing or increasing respectively as exemplified in Fig. 3D (blue and yellow curves). On the other hand, for a bounded range of Δ*E* values between the two white dashed lines (Fig. 3C), it is nonmonotonic as exemplified in Fig. 3D (red curve). The nonmonotonic dependence of MFAT on *N*_*int*_ can change depending on *N*_*pth*_ (Fig. S3). For *N*_*pth*_ = 1, MFAT shows a nonmonotonic dependence on *N*_*int*_, the same as Fig. 2C. However, the position of the minimum shifts to the right as *N*_*pth*_ is increased to 2 ∼3. The nonlinearity is then lost with a continuous increase of *N*_*pth*_. The minimum of MFAT decreases monotonically with the increase of *N*_*int*_. In order to maintain this dynamics, the number of parallel paths must also increase with *N*_*int*_ (Fig. 3E). That is, increasing *N*_*pth*_ makes the dependence of MFAT on *N*_*int*_ more monotonically decreasing. These data suggest that increasing both *N*_*int*_ and *N*_*pth*_ together accelerates the EMT process.

We have now discussed the scenarios in which there is only one path and multiple paths in parallel and found that increasing the number of the intermediate states or parallel paths could accelerate the EMT process. Here, another scenario is considered by assuming that the epithelial cell has to pass a fixed number of intermediate states to become mesenchymal. In other words, the epithelial cell must pass *N*_*ly*_ of layers to become mesenchymal (Fig. 4A). This is similar to the parallel paths scenario but has one difference. It is possible that two paths converge at one intermediate state in the layered scenario. Here, the transition rates in all steps are the same, which depends on the number of layers *N*_*ly*_.

**Fig 4.**
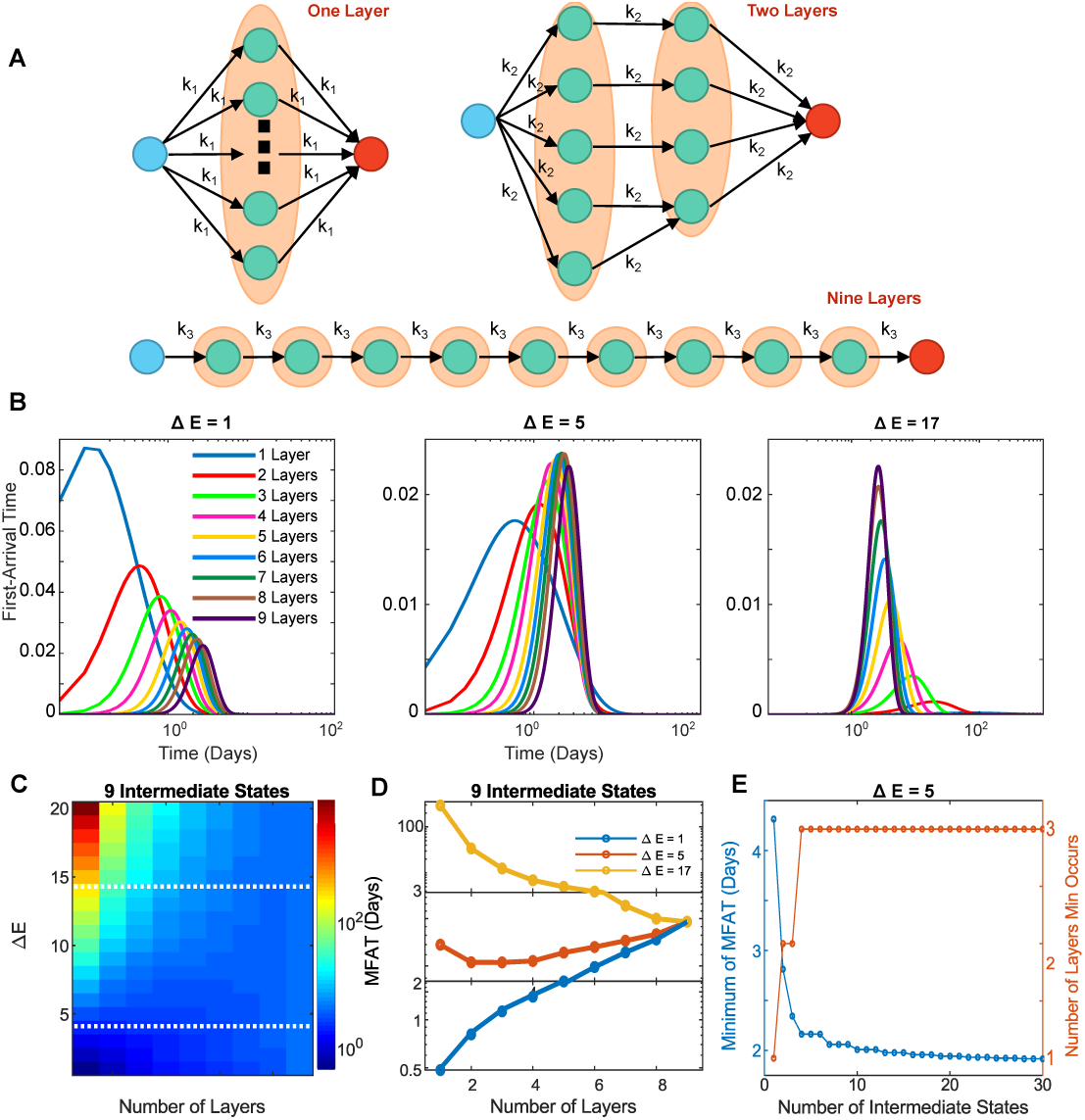
Adding transition layers can further accelerate the EMT Process. (A) Diagram of cell state transition through nine intermediate states with multiple transition layers. (B) Dependence of FAT distribution on *N*_*ly*_ with three different Δ*E* and *N*_*int*_ = 9. (C) Phase diagrams showing MFAT subjected to *N*_*ly*_ and Δ*E* with *N*_*int*_ = 9. Dashed lines represent two thresholds for monotonic and nonmonotonic dependence of MFAT on *N*_*ly*_. (D) Three typical examples show monotonic and nonmonotonic dependence of MFAT on *N*_*ly*_ according to Δ*E* value. (E) The dependence of minimum MFAT (blue curve) and the *N*_*ly*_ the minimum occurs on (red) for Δ*E* = 5.

Similar to the parallel path scenario, the first-arrival time distribution depends on Δ*E* (Fig. 4B). If Δ*E* is small, the FAT distribution shifts to the right with increase of *N*_*ly*_ (Fig. 4B, left panel), while it shifts to the left as Δ*E* becomes large (Fig. 4B, right panel). The FAT shows a nonmonotonic dependence on *N*_*ly*_ with the moderate Δ*E* (Fig. 4B, middle panel). Fig. 4C shows the MFAT in the space of *N*_*ly*_ and Δ*E* for the system with nine intermediate states. In general, a higher energy barrier gives a slower mean time arrival regardless of layer numbers. However, the trend of the MFAT versus *N*_*ly*_ depends on Δ*E*. MFAT increases with *N*_*ly*_ for small Δ*E* (below bottom white dashed line), but decreases for large Δ*E* (above top white dashed line) and shows a nonmonotonic dependence for moderate Δ*E* (between the two white dashed lines), as exemplified by the three cases in Fig. 4D. Similar to the parallel path scenario, the minimum of MFAT decreases monotonically with *N*_*int*_ as long as more layers are provided for the system with large number of *N*_*int*_ (Fig. 4E). This is also confirmed with various values of Δ*E* (Fig. 4E and Fig. S4).

Taken together, adding more parallel paths or transition layers can further accelerate the EMT process. In both scenarios, a nonmonotonic dependence of mean first arrival time on the number of paths or layers is found. The MFAT increases with the number of intermediate states after the minimal if the system only has one path, but can further decrease with multiple paths or transition layers. Thus, a combination of increasing the number of intermediate states and parallel paths (or transition layers) can always accelerate the EMT process.

### Stabilized Intermediate State Traps a Cell within its Current Phenotype

We have discussed the scenarios in which the same energy barrier is considered for each step during the EMT process. However, it is very possible that one of the intermediate states is stabilized by specific regulators [46], making the transition from this step more difficult. Here, we consider a special scenario where one of the intermediate states is more stable than the others; we call this state the “stabilized” state. As shown in Fig. 5A, our metaphoric landscape now has one well that is deeper than the others. In this stabilized state, more energy is needed for the transition to the next state. That is, the energy barrier for the transition at this step (Δ*E*_2_) is larger than the energy barrier for other steps (Δ*E*_1_). Two cases are considered here: 1) Δ*E*_2_ increases and Δ*E*_1_ decreases in order to maintain the overall energy barrier Δ*E* (Fig. 5A, “constant Δ*E* case”), and 2) Δ*E*_2_ increases as Δ*E*_1_ remains the same to change the overall energy barrier (Fig. 5A, “varying Δ*E* case”). Fig. 5B represents the diagram of the EMT process with one stabilized state, which includes two transition rates, *k*_1_ as the transition rate from the non-stabilized states, and *k*_2_ as the transition from the stabilized state.

**Fig 5.**
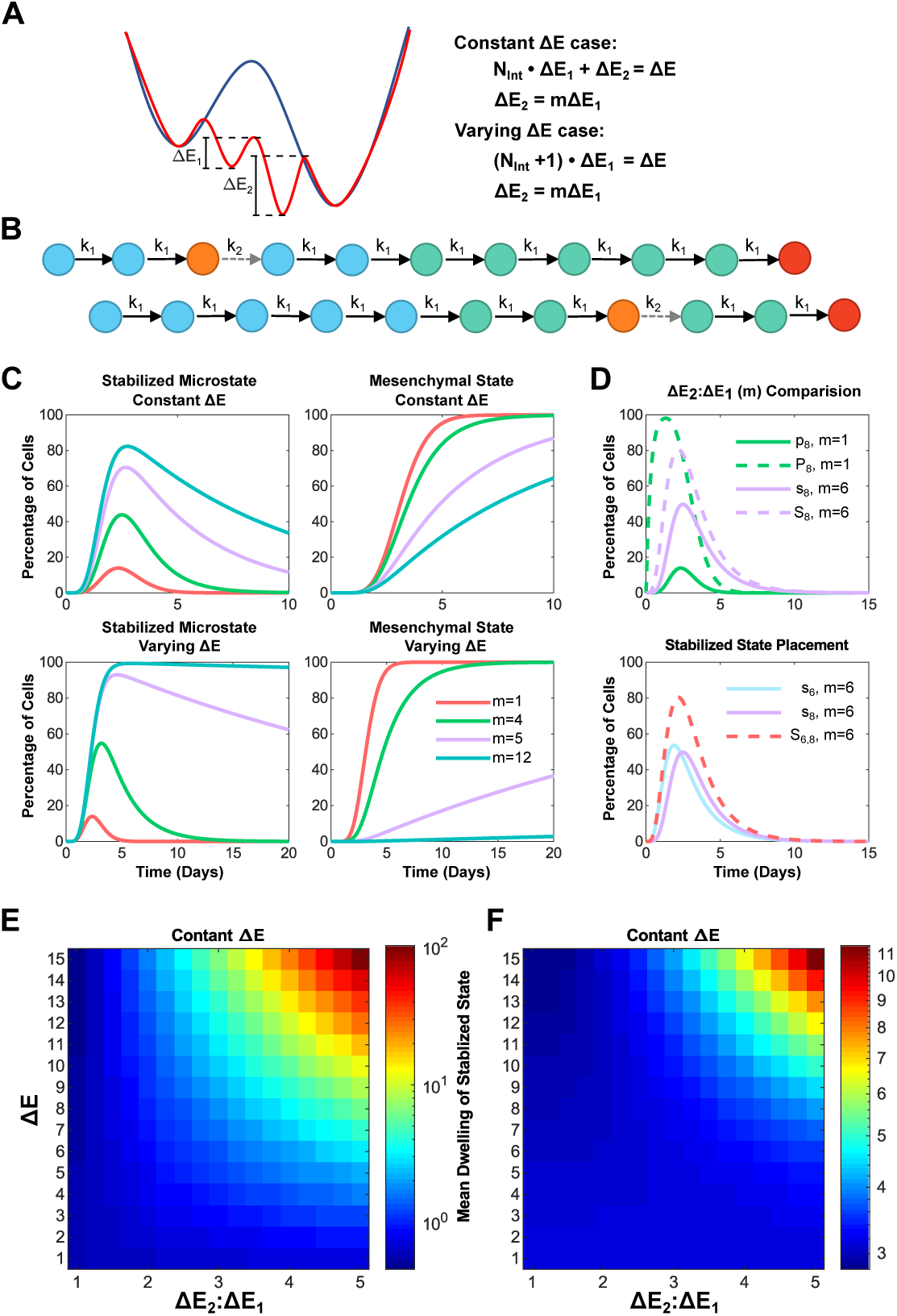
A stabilized intermediate state traps a cell within its current phenotype. (A) The metaphorical landscape of EMT with one stabilized state, which has a deeper well (Δ*E*_2_) than the others (Δ*E*_1_). Two cases were considered here: (1) the constant Δ*E* case in which Δ*E*_2_ increases and Δ*E*_1_ decreases but overall energy barrier Δ*E* remains the same, and (2) the varying Δ*E* case in which only Δ*E*_2_ increases and Δ*E*_1_ remains the same. (B) Example of a stablized state occurring within the EMT process. Orange represents the stabilized intermediate state and the gray dashed arrow represents the transition from this stabilized state (*k*_2_). (C) The distribution of cells at the stabilized state and the mesenchymal state under different energy barrier ratio, *m* (= Δ*E*_2_ : Δ*E*_1_). (D) Upper: The distribution of the cell at the stabilized microstate (solid lines) or at the corresponding macrostate (dash lines) with *m* = 1 (green) or 6 (blue). Bottom: The distribution of the cell at the stabilized microstate (solid lines) or at the corresponding macrostate with the stabilized state in 6th place (blue) or the 8th place (purple). The placement does not affect macrostate distribution. (E-F) Phase diagrams showing mean stabilized dwelling time of the stabilized state (E) and MFAT to mesenchymal (F) subjected energy barrier ratio *m* and the total energy barrier Δ*E*, in the constant Δ*E* case.

To understand the impact of a stabilized state on EMT, the distribution of cells at the stabilized state and the mesenchymal state is analyzed with different Δ*E*_2_ : Δ*E*_1_ ratios (Fig. 5C). As the ratio increases, the dwelling time inside the stabilized state grows and it takes more time to arrive to the mesenchymal state in both the constant and varying Δ*E* cases. That is, the greater the ratio, the longer the cell stays in the stabilized state, and the more time it takes to finish the EMT process. It also implies that cells stay longer in the macrostate where the stabilized intermediate state belongs (Fig. 5D, upper panel) and that the placement of the stabilized intermediate state within this macrostate does not affect the macrostate behavior (Fig. 5D, bottom panel). The dependence of the mean dwelling time within the stabilized state and the MFAT is further analyzed. As shown in Fig. 5E-F, with increase of either the ratio Δ*E*_2_ : Δ*E*_1_ or the overall energy barrier, the mean dwelling time within the stabilized state and the MFAT increase. The same dependence is found for the varying Δ*E* case (Fig. **??**) but the scale of changes significantly increases. Another difference between two cases is the reverse dependence of MFAT on Δ*E*_2_ : Δ*E*_1_ as shown in (Fig. **??**) when on the Δ*E* is small. Taken together, a greater overall energy barrier or greater intermediate state energy barrier ratio will cause for a slower transition into the mesenchymal state.

## Discussion

Multiple partial EMT states have been proposed and verified to exist in the EMT process, and the number of states involved depends on the cell line. Here, we focused on the dynamics and functions of EMT plasticity. We provided microstate and macrostate concepts for EMT. Based on the parameter fitting of a Markov model of EMT with the experimental data, we proposed a statistical mechanism for EMT in which many unobservable microstates exist in one of the observable macrostates. That is, EMT is a process with many microstates, which can be mapped into different observable macrostates. The microstates may encompass other dimensions of the cell that are coupled to the EMT process, such as the cell cycle, stemness, and metabolism. How the number of microstates is determined is not clear but largely depends on the cell lines, which may be revealed with analysis of sing-cell time-course data [47].

Here, we performed a systematical analysis on how the number of intermediate states changes cell transformation. We considered several scenarios including single transition path, parallel transition, and layered transition. We found that in the single transition path scenario, increasing the number of intermediate states can accelerate EMT, but too many intermediate states also have a potential of decelerating EMT, thus showing a nonmonotonic behavior. However, adding parallel paths or transition layers can further accelerate the EMT process, especially for a system with large number of intermediate states.

Our results suggest that the number of intermediate EMT states may function as an indicator for the malignancy of the cancer. That is, the more intermediate EMT states in one cancer, the more chance it will metastasize. Thus, the malignancy level can be quantified by the number of potential intermediate EMT states. It is estimated that metastasis causes 90% of cancer deaths [48]. If we can stop or slow down the transition, we have a great chance of treating cancer. Our results suggest one potential strategy targeting on the EMT spectrum, i.e., reducing the number of the potential partial EMT states.

Lastly, the existence of a stabilized state within the EMT process causes a slower transition. It is reported that cells in the partial EMT state contribute more to cancer metastasis than the mesenchymal state [49]. Further more, some partial EMT states can be stabilized by the cancer cells [50]. Our simulation shows that stabilizing one intermediate state will trap the cells there for a much longer time period, slowing down full EMT. The multiple intermediate states emerge due to various interlinked positive feedback loops formed in the regulatory network of EMT, which has been demonstrated in many previous works that combined mathematical modeling and quantitative experiments [5–11, 45]. It also has been shown that removal of counteracting determinants traps cell in the rare myeloid transition state [51]. Similarly, if we can tune the strength of the feedback loops that are responsible for the generation of these partial EMT states, the partial EMT states will destabilize and even disband. This will make the first step of metastasis mediated by EMT or partial EMT more difficult and thus give us more time to treat the cancer.

Here in this work, we did not consider the molecular mechanism of the intermediate states but focused on the how the number of the intermediate states affect the EMT process. Further work is needed to verify the prediction derived here and the detailed molecular mechanism needs to be determined for the potential therapeutic targets. Our results suggest that a small perturbation of the signaling pathway could change the EMT process significantly and thus EMT-related diseases, such as cancer metastasis and renal fibrosis. Our previous work on renal fibrosis suggests that knockout the EMT genes might not be the optimal treatment design for acute kidney injury and long-term firbrosis [41]. Thus, future works can search control strategies to slow down or accelerate EMT with dynamic perturbation of the signaling pathway toward optimal treatment of cancer or fibrosis.

The accelerating function of intermediate states on EMT is analogous to the phenomena in protein folding [52]. It is becoming increasingly apparent that many other cellular processes have multiple intermediate states instead of just one binary or continuous process. For example, stem cell differentiation consists of many intermediate states [53–55]. Thus, it will be exciting to analyze one biological process in different perspectives, including statistical mechanics, mathematical modeling, and single-cell analysis.

## Author Contributions

X-J.T. and L.C. conceived the study. X-J.T., L.C. H.G., and J.M-A. designed the study. X-J.T., H.G., and J.M-A. performed model studies. X-J.T. and H.G. analyzed the data and wrote the manuscript with inputs from all authors.

## Acknowledgments

This project was supported by the ASU School of Biological and Health Systems Engineering and NSF grant (EF-1921412)(to X-J.T.). H Goetz and J Melendez-Alvarez were also supported by the Arizona State University Dean’s Fellowship.

## Supporting information

## Supplementary Figures

**Fig S1.**
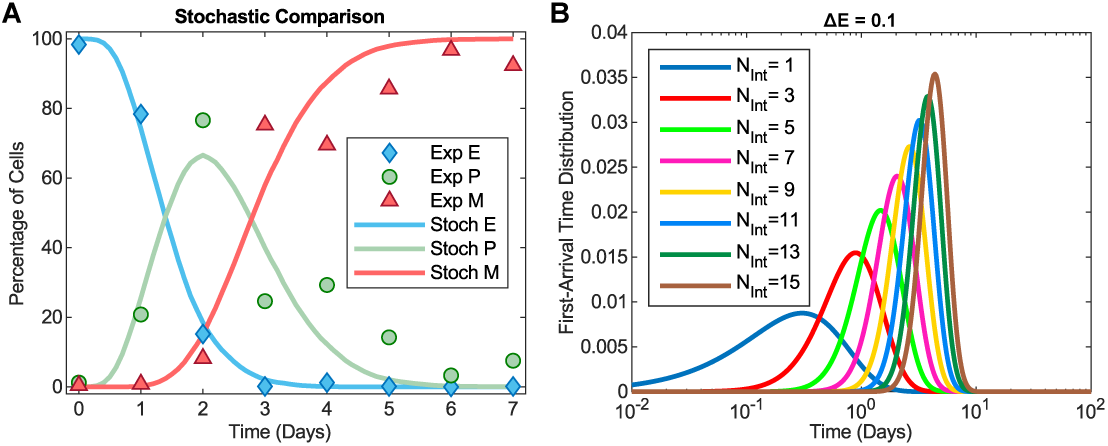
(A) The dynamics of the EMT process with the stochastic model overlaid with experimental data [6]. The simulation data is sampled with 10,000 cells from the stochastic model. (B) The FAT distribution to the mesenchymal state with different numbers of intermediate states and Δ*E* = 0.1.

**Fig S2.**
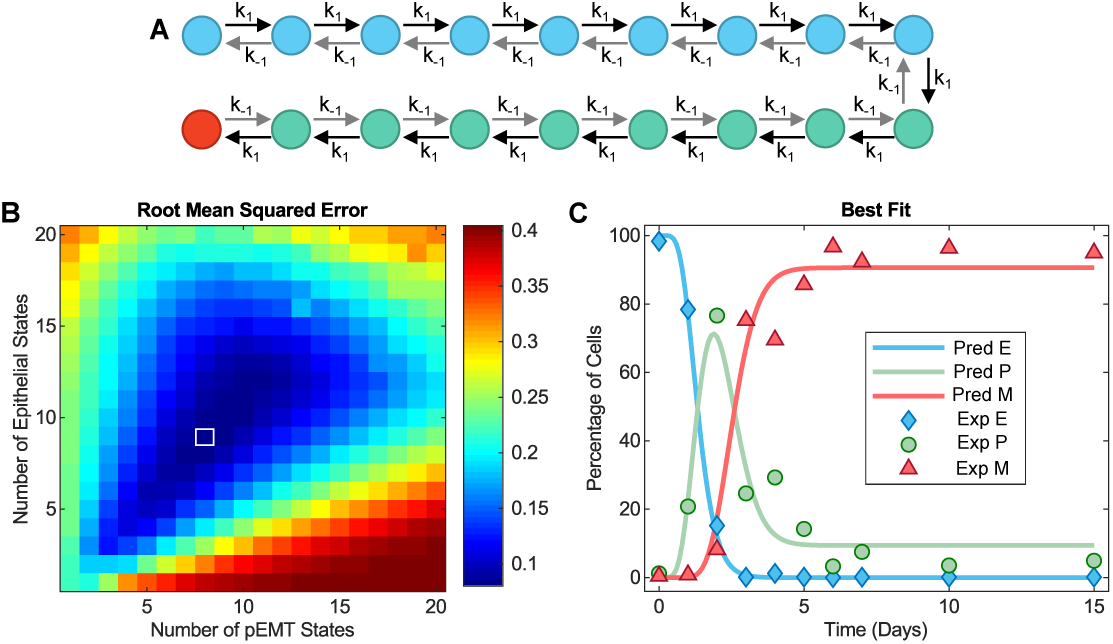
Fitting the experimental data with a model that considers reversible cell state transitions suggests more microstates during EMT.. (A) EMT progression through a continuum of reversible intermediate states. (B) The fitting score, root mean squared error (RMSE), for the model with reversibility at the space of *N*_*E*_ and *N*_*P*_ gives the best fit at *N*_*E*_ = 9 and *N*_*P*_ = 8 (RMSE=0.0064, *k*_1_ = 7.1258 and *k*_*-*1_ = 0.6694). (C) The dynamics of the EMT process with the best-fitted model of reversible intermediate states overlaid with experimental data from Ref. [6].

**Fig S3.**
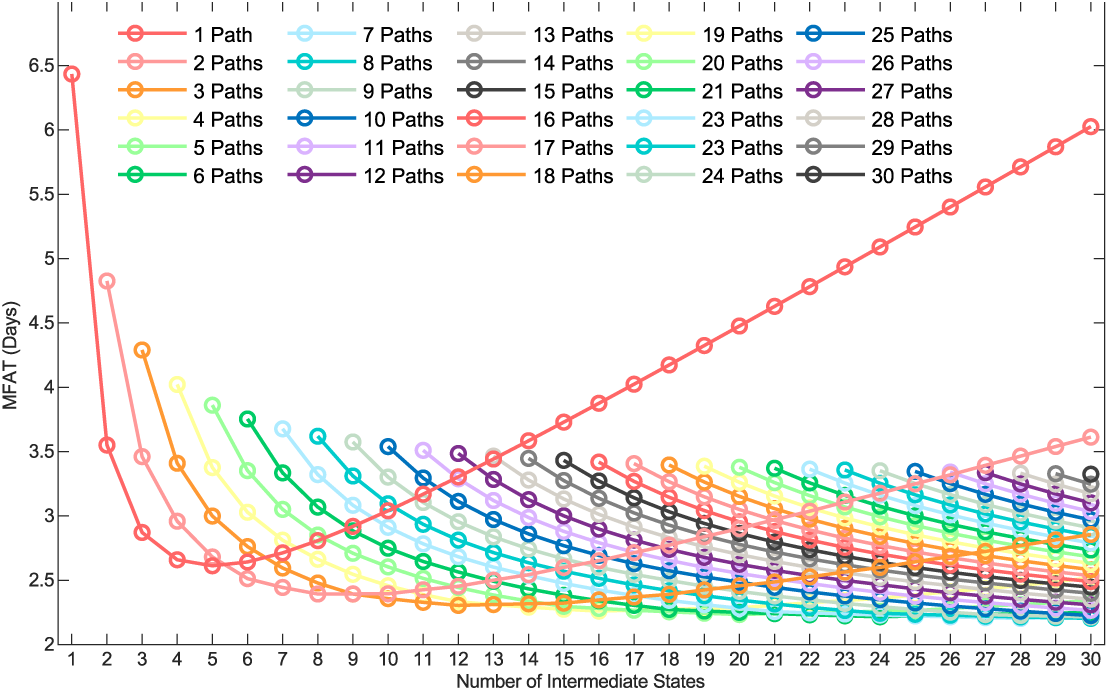
Adding parallel paths changes the dependence of MFAT to the mesenchymal state on the number of intermediate states. MFAT as a function of *N*_*int*_ under various *N*_*pth*_ with Δ*E* = 6.

**Fig S4.**
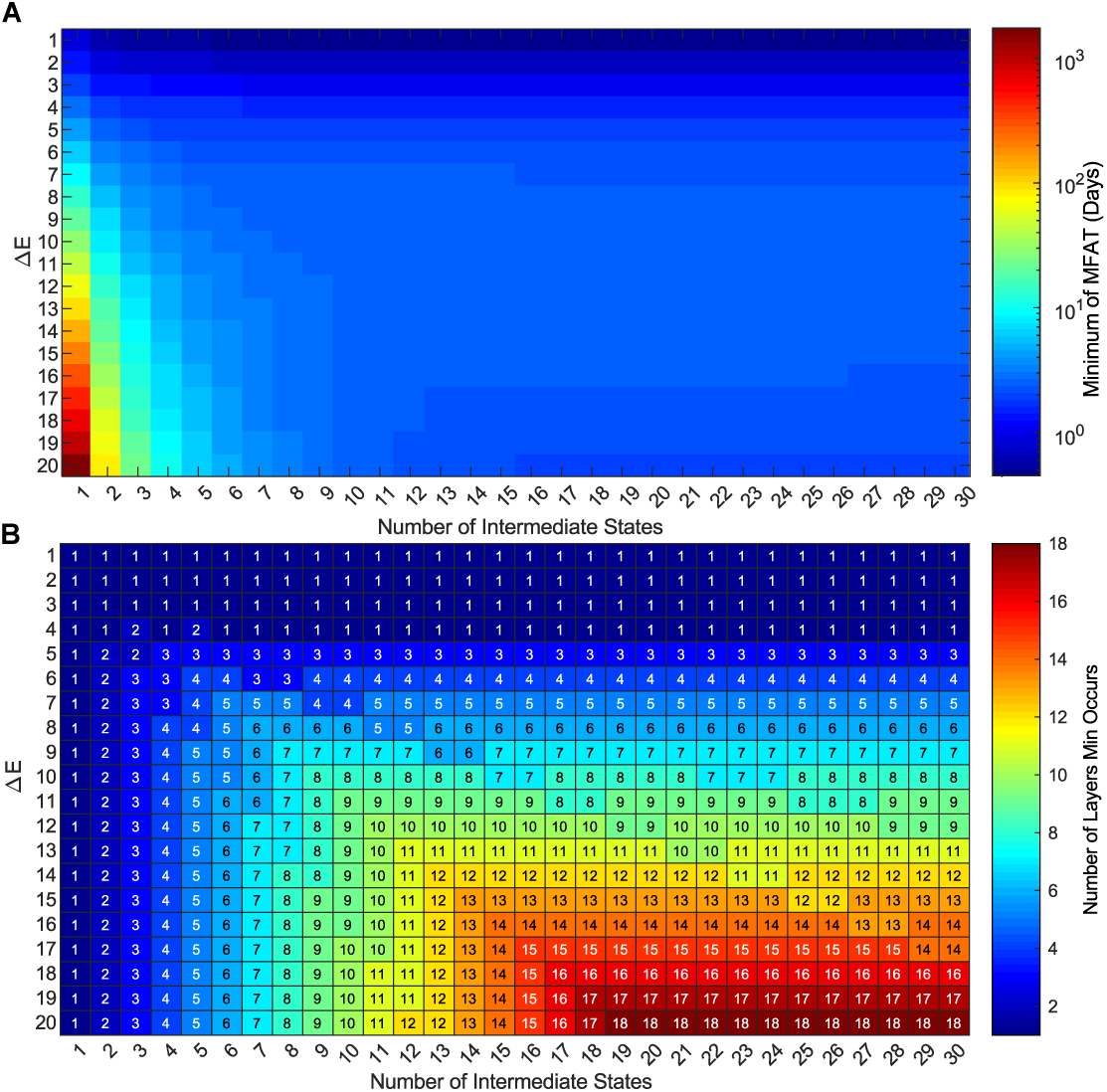
Adding transition layers changes the dependence of MFAT to the mesenchymal state on the number of intermediate states. Minimum MFAT (A) and the corresponding number of layers in the space of number of intermediates states *N*_*int*_ and Δ*E*.

**Fig S5.**
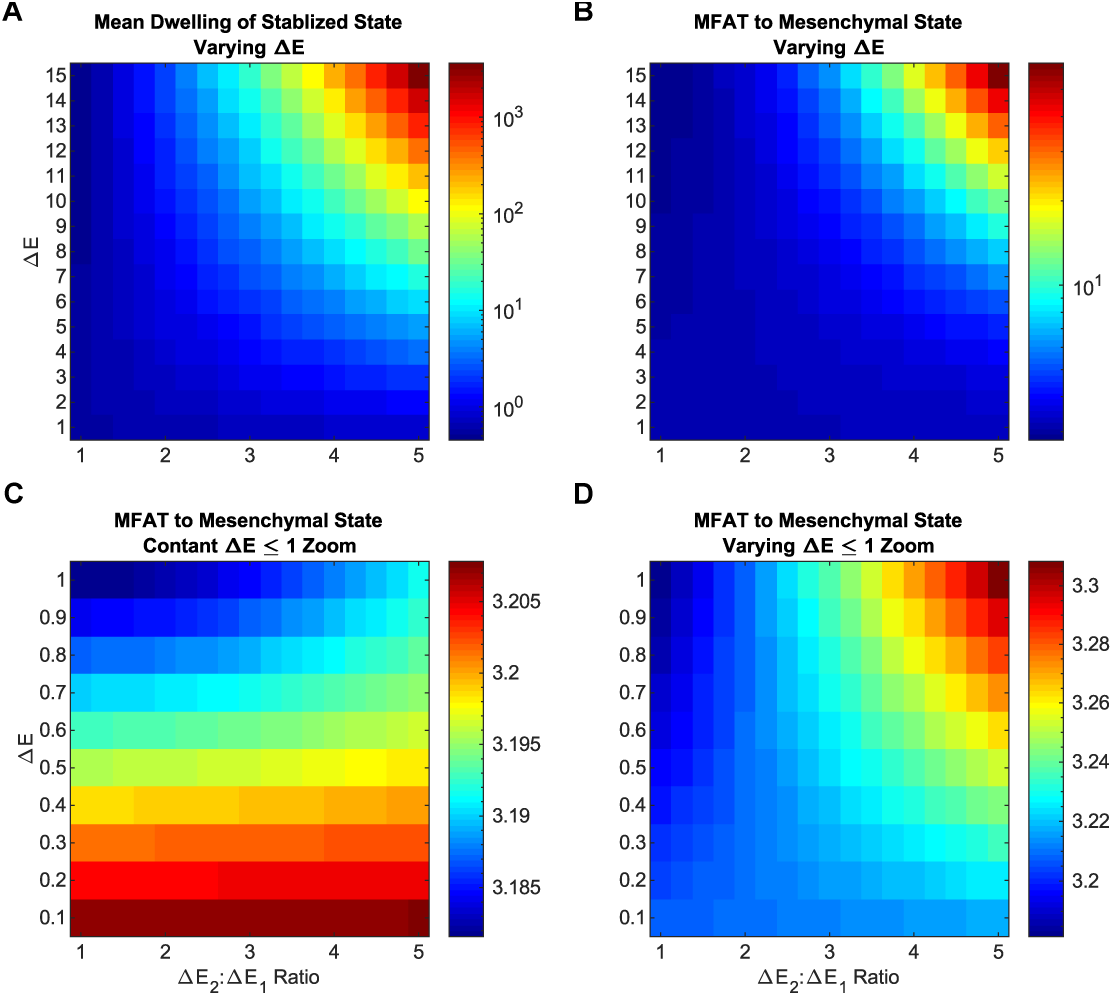
Comparison of two mechanisms of stabilizing one intermediate state on EMT dynamics. (A-B) Phase diagram showing mean stabilized dwelling time of the stabilized state (A) and MFAT to the mesenchymal state (B) with energy barrier ratio Δ*E*_1_ : Δ*E*_2_, and total energy barrier Δ*E* in the varying Δ*E* case. (C-D) MFAT for Δ*E* ≤1 with energy barrier ratio Δ*E*_1_ : Δ*E*_2_ and total energy barrier Δ*E* in the constant Δ*E* case (C) and the varying Δ*E* case (D).

## Supplementary

### Transition Rate and Energy Barrier

In this work, we assume that the total energy barrier one epithelial cell has to cross to be transitioned to the mesenchymal state is Δ*E*. If there is no intermediate state, then following the Arrhenius equation, the transition rate of EMT is 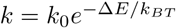, where the value of *k*_0_ is a pre-exponential factor, which determines the time scale of the system. For simplicity, we define the energy unit as *k*_*BT*_, thus we can simplify the equation to *k* = *k*_0_*e*^*-*Δ*E*^. Suppose there are *N*_*int*_ intermediate states between the epithelial and mesenchymal states and that the energy is divided into *N*_*int*_ + 1 steps; the transition rate for step *i* is 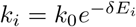 where *δE*_*i*_ is the energy barrier for this step and Σ *δE*_*i*_ = Δ*E*. We considered two scenarios:

1. The energy is divided evenly into all the steps, then the transition rate for each step is same as 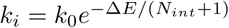.
2. If there is one stabilized intermediated state, where the energy barrier is *δE*_*s*_ = *mδE*_*i*_. We considered two cases, a) *δE*_*s*_ + Σ_*i*≠*s*_ *δE*_*i*_ = Δ*E*. The transition rate is 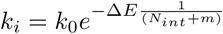 for the transition from the regular intermediate states, and 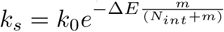 for the transition from the stabilized intermediate state; b) The transition rate is 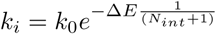 for the transition from the regular intermediate states (same as scenario 1), and 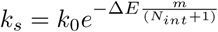 for the transition from the stabilized intermediate state;

### Estimation of *k*_0_ and the Energy Barrier Δ*E*

The total energy barrier from epithelial to mesenchymal, Δ*E*, was estimated by the following equations:

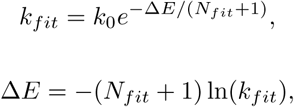

where *N*_*fit*_ = 9 is the number of intermediate states and the transition rate *k*_*fit*_ = 3.4261 from the best fitting of the experimental data in Fig. 1.

### Parameter Fitting

#### Based on One Intermediate State and an Irreversible EMT Process

First, we considered one intermediate state in the model to fit the parameters. We assumed EMT to be an irreversible process under high dose of inducer TGF-*β*. 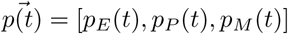 are the probability of cells in each state during the process of EMT. Thus, the equation governing the dynamics of 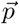 is

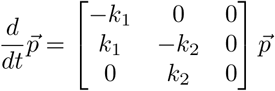

where, with *k*_1_ as the transition rate from the epithelial state to the pEMT state, and *k*_2_ is the transition rate from the pEMT state to the mesenchymal state.

The equations are solved and the solution is:

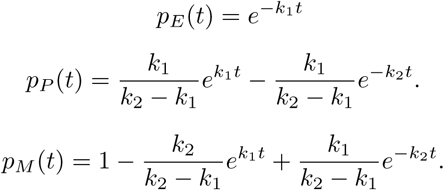

#### Based on a Flexible Number of Intermediate States and an Irreversible EMT Process

The above model does not describe the experimental data well (Fig. 1C). Thus, we extended the model by considering *N*_*int*_ + 1 microstates; this is based on an assumption that each macrostate, including the epithelial state and pEMT state, consist of multiple microstates. Suppose that EMT is an irreversible process where the cells transition independently from one microstate to the next at the same rate *k*. Based on the assumption that the Δ*E* is evenly divided into these steps, the following equations can be used to represent the dynamics of the probability of the microstates:

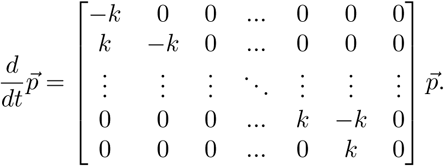

The solution of this equation is:

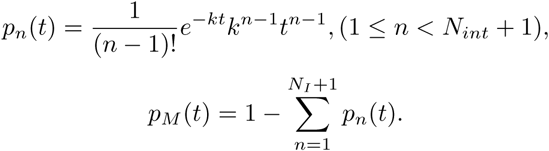

This model is used for fitting the experimental data as shown in Fig. 1F.

#### Based on a Flexible Number of Intermediate States and a Reversible EMT Process

EMT has some probability to be reversible even under high dose of inducer TGF-*β*, which may result from the noise in the cell or other stochasticity. We also considered a case where the EMT process is reversible with the cells transitioning independently between one microstate and the next at the forward rate *k*_1_ and backward rate *k*_*-*1_. The following equations can be used to represent the dynamics of the probability of the microstates:

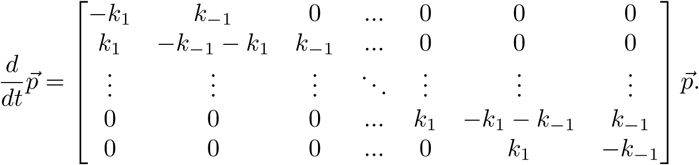

This model was also used for fitting the experimental data as shown in Fig. S2C.

### First Arrival Time

The first arrival time (FAT) distribution is used to quantify the time cells take to be transited into the mesenchymal state in various cases. In the case of an irreversible EMT process with microstates, the FAT distribution is 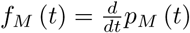 with normalization 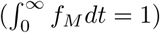. Then,

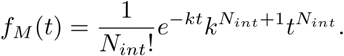

The corresponding mean first arrival time (MFAT) can then be found with 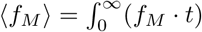, thus

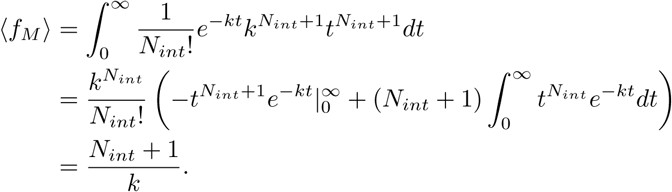

### The Cases with One Stabilized State

For any case that has one stabilized state at *S*, such that 1 *< S < N*_*int*_ + 1, the pattern goes as follows:

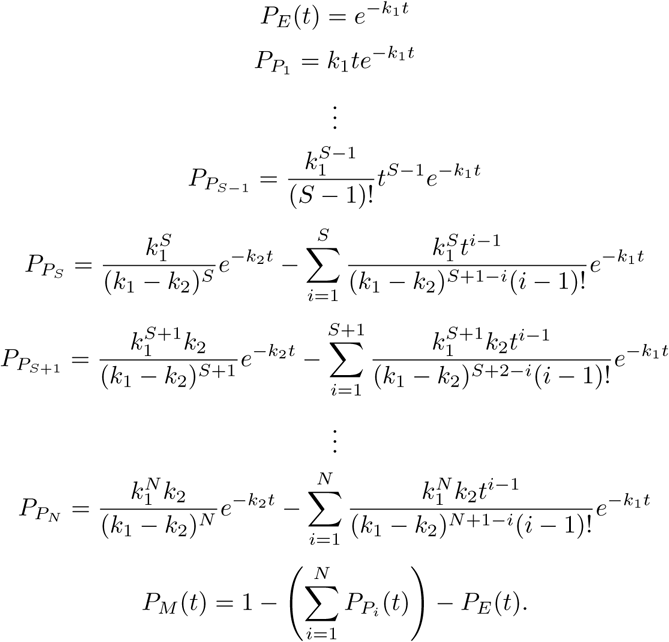

Here, the *k*_1_ denotes the transition rate from the non-stabilized state and *k*_2_ the transition from the stabilized state. In order for the state to be stabilized as described, it must be true that *k*_2_ *< k*_1_.

### Stochastic Model

To simulate the stochastically of the cell state transitions during EMT at the single-cell level, we also developed a stochastic model using the Gillespie algorithm.

1. Define the system according to the cell state transitions during EMT, using matrix *S* is the cell state the transition from, matrix *P* is the cell state the transition to, vector *K* is the transition rates. For example, for the system with only one path and *N*_*int*_ intermediate states, *S* and *P* have *N*_*int*_ + 2 columns and *N*_*int*_ + 1 rows while *K* has one column and *N*_*int*_ + 1 rows, with the pattern:

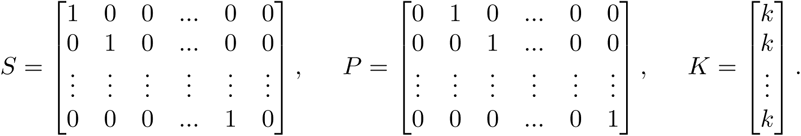
2. Initialize the system. Time *t*_0_, the cell state *x*_0_ = [1; 0; …; 0]*asonecolumnvector*.
3. Determine the rate of each cell transition *a* = *Sx*_0_*K*, and *a*_0_ = Σ *a*.
4. Generate two uniformly distributed random number *rn*_1_ and *rn*_2_ between [0 1].
5. Determine the time it takes for the cell state transition *dt* = *ln*(1*/rn*_1_)*/a*_0_,
6. Determine the step cell state transition occurs *r* which satisfies 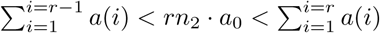.
7. Update the system *x*_0_ = *x*_0_ - *S*(*r*, :) + *P* (*r*, :) and *t* = *t* + *dt*.
8. Repeat step 3 - 7 until *t* is more than maximum time *T*_*max*_.

This stochastic model was used in Fig. 1G-H. The parameters are taken from the best fit in Fig. 1E-F.

